# Measuring excitation-inhibition balance through spectral components of local field potentials

**DOI:** 10.1101/2024.01.24.577086

**Authors:** Geoffrey W Diehl, A David Redish

## Abstract

The balance between excitation and inhibition is critical to brain functioning, and dysregulation of this balance is a hallmark of numerous psychiatric conditions. Measuring this excitation-inhibition (E:I) balance *in vivo* has remained difficult, but theoretical models have proposed that characteristics of local field potentials (LFP) may provide an accurate proxy. To establish a conclusive link between LFP and E:I balance, we recorded single units and LFP from the prefrontal cortex (mPFC) of rats during decision making. Dynamic measures of synaptic coupling strength facilitated direct quantification of E:I balance and revealed a strong inverse relationship to broadband spectral power of LFP. These results provide a critical link between LFP and underlying network properties, opening the door for non-invasive recordings to measure E:I balance in clinical settings.

## Main Text

Network communication within the brain is critical for representing, relaying, and processing information (*1, 2*). In particular, neural functioning relies on an exquisite balance between excitatory communication and inhibitory control (*3-7*). As evidence for the importance of this balance between excitation (E) and inhibition (I), multiple psychiatric conditions have been hypothesized to arise from a dysregulation of the E:I ratio (*8-12*). However, obtaining quantitative measures of this network communication is often difficult, particularly in humans, and particularly in clinical settings where the invasive techniques necessary to do so are unavailable in all but the most extreme cases. Recent work suggesting that electroencephalogram (EEG) and local field potential (LFP) signals arise as an emergent property of excitatory and inhibitory post-synaptic potentials has put forward the hypothesis that directed analysis of EEG and LFP could provide a viable proxy for changes in E:I balance. Specifically, these models have proposed that the aperiodic component of spectral power (commonly the 1/f component) provides an estimate of the real-time balance between excitation and inhibition (*13-15*). As yet, however, this hypothesis has not been directly tested in the complex conditions of dynamic real-world behavior. To directly test this hypothesis under real world behavior, we leveraged a unique data set of largescale physiology recordings taken from silicon probes in the medial prefrontal cortex (mPFC) of rats running a complex, naturalistic decision-making behavior (Restaurant Row; RRow) (*16*). We extracted direct measures of excitatory and inhibitory synaptic coupling strength and simultaneous LFP signals, and found that the proposed relationship between the LFP aperiodic component and the E:I balance did not hold true. Instead, E:I balance was strongly related to broadband spectral power.

Across 8 rats and 104 decision making sessions we recorded 9344 channels of LFP and 3017 single units (*16*), which provided a uniquely rich data set in which we could evaluate a total of 107418 neuron-to-neuron interactions, from which we could directly measure how excitatory and inhibitory communication changed during dynamic behaviors. We computed high-resolution spike cross-correlation histograms (CCHs) for each pair of simultaneously recorded cells (Fig 1A-C, S1A), a method that is commonly used to identify and characterize cell pairs with a monosynaptic connection (*17, 18*). We identified 3142 significant CCHs and over 1500 instances of putative synaptic connections (Fig 1D-G, S1B-C), with both excitatory and inhibitory connections appearing in roughly equal quantity (782 excitatory and 766 inhibitory connections). The CCH analysis also revealed 864 instances of synchronous co-spiking between pairs of mPFC cells (Fig S2), a phenomenon that has been associated with long-range coordination across the brain and with proper functioning of NMDA receptors (*19-24*). Although these putative monosynaptically coupled pairs represented a very small proportion of the overall candidate cell pairs (1.5%), the scale of our recording data meant that we identified a sufficient number of coupled pairs to effectively evaluate their relationship to ongoing LFP.

**Figure 1:**
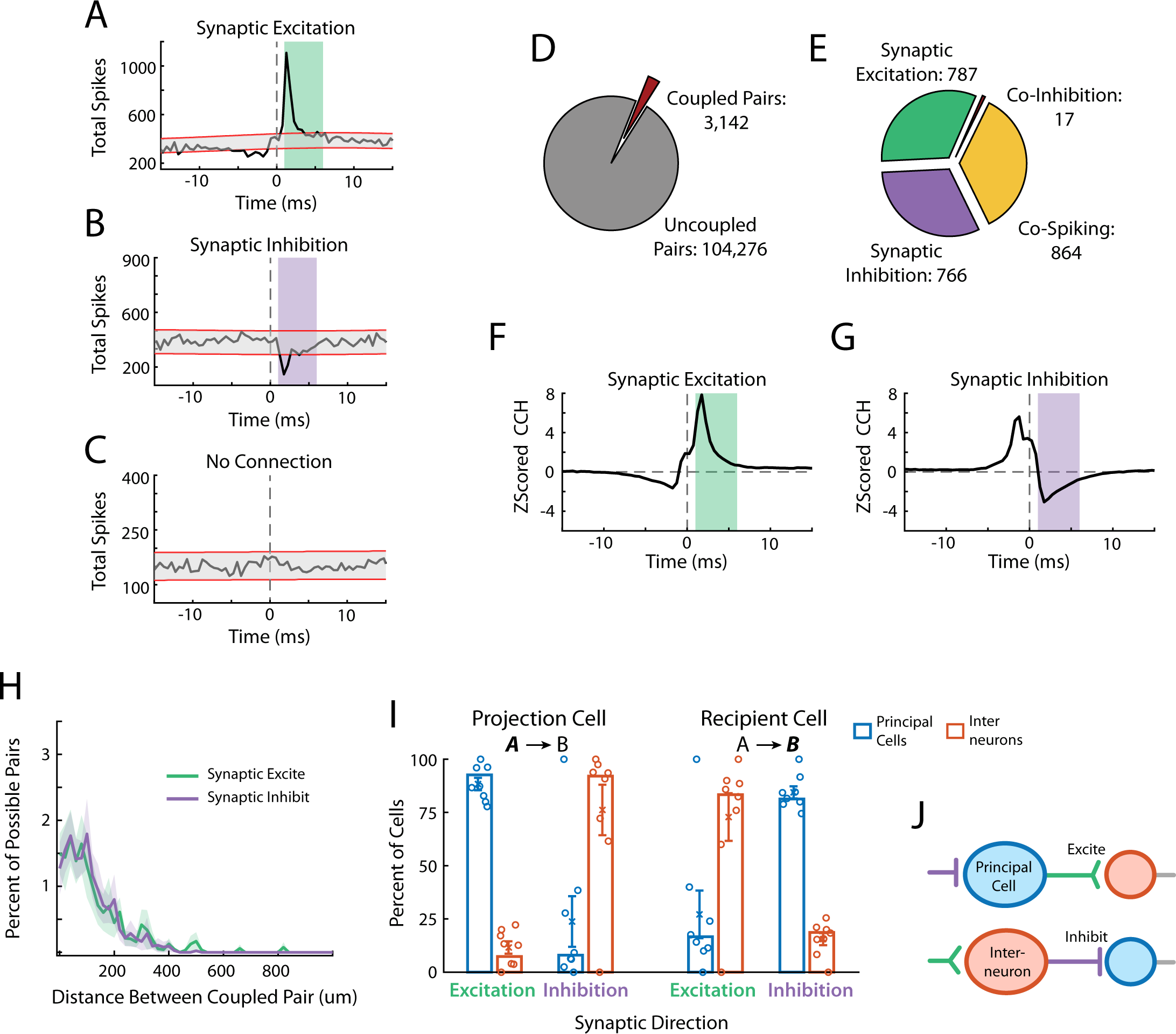
Coupling between mPFC cell pairs. (A-C) Cross correlation histograms (CCHs) of three pairs of simultaneously recorded mPFC cells. Analysis identified an excitatory connection (A; *R535-2019-07-21 SiC1_33 ◊ SiC1_37*), inhibitory connection (B; *R535-2019-07-21 SiC1_07 ◊ SiC1_13*), or no detectable connection (C; *R535-2019-07-21 SiC1_33 ◊ SiC1_25*). Confidence bounds around the expected CCH rate are shown in red and the time windows for detecting synaptic connections are shown as green or purple shading. D) The number of all possible cell pairs in which CCH analysis identified a significant interaction. E) Number of cells pairs identified in each possible category. Note that Co-Spiking pairs count twice in D but only once in E (both *candidate pairs* were “coupled” but this represented only a single interaction), and that a given cell pair could be counted in multiple categories. (F,G) Average z-scored CCH responses for those cell pairs with identified synaptic connections. H) The frequency of identifying synaptic couplings as a function of the anatomical distance between the cell pairs. I) Percent of excitatory or inhibitory synaptically coupled cells that were identified as “principal cells” or “interneurons” based on waveform and firing rate. Left and right bars independently quantify the projection cell or the recipient cell in the coupled pair. Proportions across the full data set are shown as bars. Proportions from each rat are shown as circles and error bars denote the mean ± SEM across rats J) Our data are consistent with a ping-pong style architecture in which information flows alternatively between excitatory and inhibitory network components.

Notably, our data set, and the identification of excitatory and inhibitory connections within it, provided valuable insight into underlying properties of network architecture within the mPFC (Fig S3, S4). The use of silicon probes allowed us to identify the precise anatomical location of each recorded cell (*16, 25*) and to directly quantify the anatomical relationship between coupled cells. In line with cortical network models (*20, 26-28*), the proportion of synaptic connections we identified were inversely related to the distance between the cells in a pair (Pearson’s correlation distance vs proportion: r=-0.75, p<0.001), with virtually no couplings occurring at distances greater than 500 µm (Fig 1H, S2D). Interestingly though, this distance relationship held for both excitatory and inhibitory connections (Excite: r=-0.74, p<0.001; Inhibit: r=-0.74, p<0.001), running counter to theories of network inhibition acting broadly and in contrast to localized excitation (*6, 26, 29*). Instead, our data are in line with mPFC networks operating within a columnar organization, where cells preferentially communicate within a prescribed local region (*30, 31*). Indeed, anatomical work has identified cortical columns within the rodent mPFC to span 250– 750µm (*32*), consistent with our data.

Additionally, putative principal cells and interneurons have long been identified from overall firing rate and extracellular waveform characteristics (*33, 34*). While principal cells and interneurons classified based on these metrics are often assumed to reflect excitatory and inhibitory cells respectively, our CCH analysis allowed for direct verification of this fundamental assumption in the mPFC *in vivo*, at scale, during freely-moving behavior. Notably, we found over 90% correspondence between these two analysis methods, with principal cells delivering almost all synaptic excitation, and interneurons responsible for virtually all synaptic inhibition (Fig 1I). Furthermore, this analysis revealed that principal cells and interneurons typically received opposing synaptic input (inhibition and excitation respectively) (Fig 1I, S4C), suggesting that the mPFC network can be largely characterized according to a ping-pong style architecture (*26*) in which information is preferentially relayed back and forth between excitatory and inhibitory populations (Fig 1J).

To directly test prior theories relating the LFP aperiodic spectrum to E:I balance (*13, 14*), we combined our CCH-based identification of excitatory and inhibitory synaptic coupling with a dynamic quantification of the exponential decay rate of the LFP spectrum throughout the behavioral session (Fig 2A-C). Using 512ms bins, the exponent slope of the LFP spectral power from 1Hz to 150Hz varied dynamically throughout the session (Fig S5A-G), while also remaining highly consistent across LFP channels, behavioral session, and individual rats (Fig S5H-J, S6).

**Figure 2:**
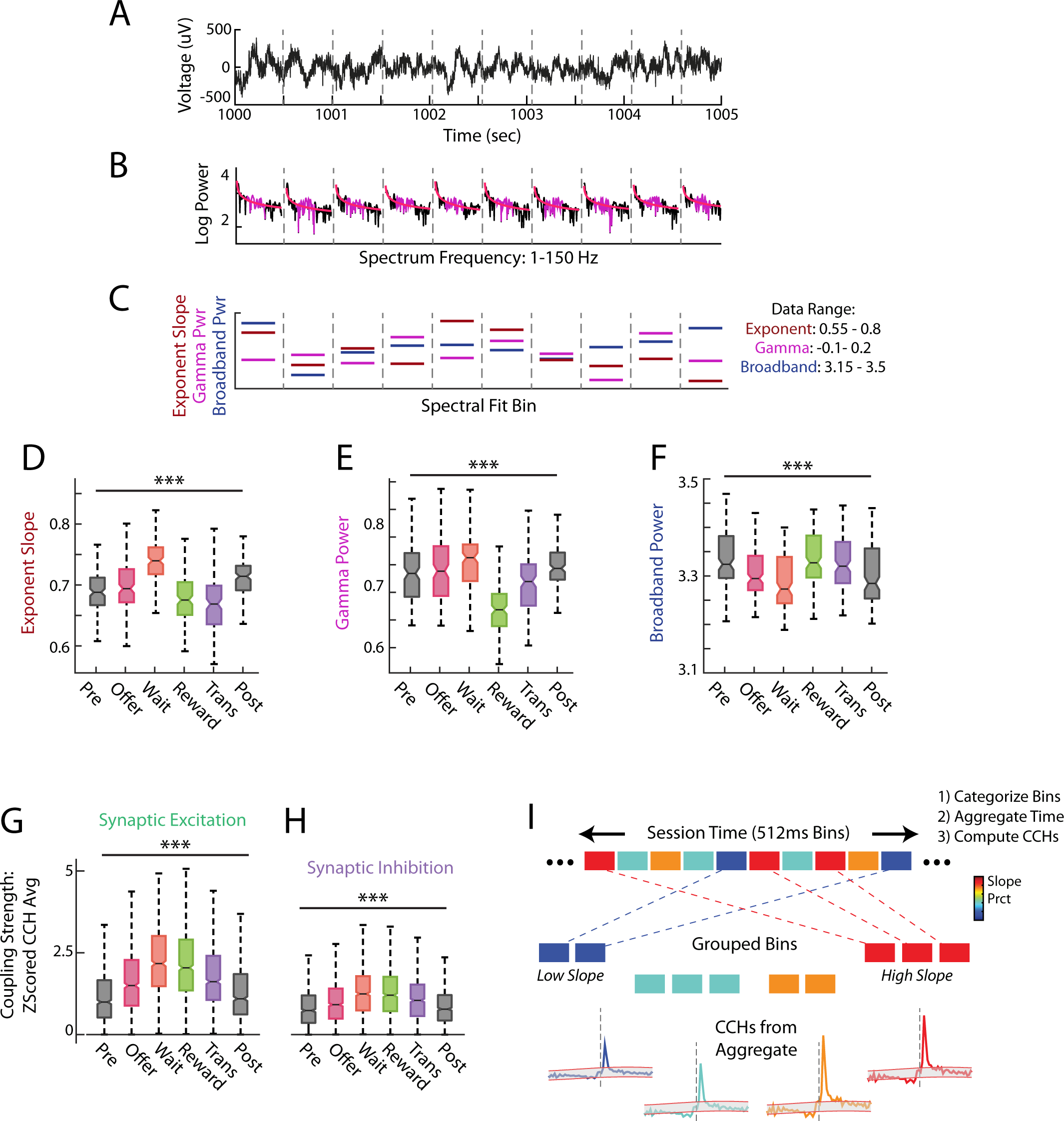
Quantification of LFP characteristics. A) Example trace of raw LFP recording (*R542-2019-07-21 SiC1_14*). Vertical dashed lines indicate the bounds between 512ms spectral analysis windows. B) A series of power spectra calculated from each window of the example LFP. Each spectrum is computed from 1-150Hz with frequency on the x-axis resetting in each window. The exponential fit is shown in red and the gamma range (35-95Hz) is colored magenta. C) Quantification of the exponential fit slope, net gamma power above the aperiodic, and average broadband power in each spectral window. Note that each metric is shown on its own scale for illustration purposes and the min-max ranges are noted right. (D-F) Quantification of LFP components across different phases of the Restaurant Row task, or the pre/post task epochs. (G,H) Quantification of synaptic coupling strength from CCHs computed during different phases of the task or pre/post epochs. I) Schematic outlining the method for relating CCH responses to LFP characteristics.

To explicitly measure the dynamic nature of the LFP aperiodic slope, we segregated our recording data according to rats’ progression through the Restaurant Row task. Past work has revealed that different components of RRow engage distinct cognitive processes and neural dynamics (*16, 35, 36*). If, as proposed, the LFP aperiodic slope provides a lens into these underlying dynamics, we would expect the slope to vary systematically across our behavioral task. Indeed, this was what we found (Fig 2D-F; ANOVA across behavior; Exp. Slope: F(5)=47.3, p<0.001; Gamma: F(5)=55.3, p<0.001; Broadband: F(5)=5.8, p<0.001). The LFP aperiodic slope was not uniform across task phases but was instead higher (steeper spectrum) during the “Wait Zone” when rats waited out a temporal delay, and was lower (shallower spectrum) in the “Reward Zone” when rats received their food reward.

Notably, synaptic coupling strength also varied systematically across the task (Fig 2G,H, S7; ANOVA; Excite: F(5)=162.5, p<0.001; Inhibit: F(5)=59.8, p<0.001) with stronger communication during periods when rats were anticipating or consuming their earned reward (Wait Zone and Reward Zone). Thus, our unique experimental paradigm and *in vivo* measures of synaptic coupling strength and ongoing LFP provide a critical opportunity to directly probe the underlying relationship between E:I balance and emergent LFP characteristics during dynamic, real-world conditions.

While our analysis of LFP characteristics allowed for the assessment of dynamics at a time resolution of 512ms, analysis of cell coupling via CCHs requires a larger amount of spiking data than is available in these short time windows. To address this limitation, we developed a novel analysis that allowed us to effectively measure excitatory and inhibitory synaptic strength at varying levels of aperiodic slope (Fig 2I). Briefly, aperiodic slopes from each 512ms time window were grouped into 20 categories, according to their percentiles throughout the session. Time windows from each percentile category were then aggregated together yielding about 3 minutes of total recording time corresponding to a given level of aperiodic slope. CCHs were computed from spiking in the aggregated 3 minutes, excitatory and inhibitory strength was extracted for each cell pair in each LFP percentile category, and the dynamic E:I balance was computed as the ratio of average synaptic excitation to inhibition. This novel approach to *in vivo* recordings and the richness of our data set thus enabled a direct experimental measure of the local E:I balance within the mPFC as a function of corresponding dynamic characteristics of the ongoing local field potential.

Past work has theorized that the aperiodic spectrum of local field potentials arises as an emergent property of the balanced integration of excitatory and inhibitory synaptic input (*13, 15*). Therefore, it has been hypothesized that measures of the exponential slope of this aperiodicity (in the 100Hz range) is inversely proportional to the ratio of excitation to inhibition 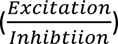 (Fig 3A) with shallower spectrum (smaller magnitude slopes) corresponding to more excitation and less inhibition. As a direct test of this theory, we correlated the E:I ratio computed from CCHs in each 3-minute aggregate of session time with the corresponding average aperiodic slope (Fig 3B). Quite surprisingly, our data did not support the proposed model relating aperiodic slope to E:I ratio. If fact, we observed a mild *positive* correlation between LFP and underlying spiking dynamics (Mann-Whitney U test vs shuffle; Exp. Slope: z=2.02, p=0.04, d’=0.26), in stark opposition to our expectation. Notably, these same findings held if we used overall spiking rates of principal cells and interneurons as our proxy for E:I balance in the local network (Fig S8). Thus, not only do our experimental data fail to lend direct experimental support to the original hypothesis, they argue for a need to fundamentally re-evaluate the relationship between LFP and E:I balance.

**Figure 3:**
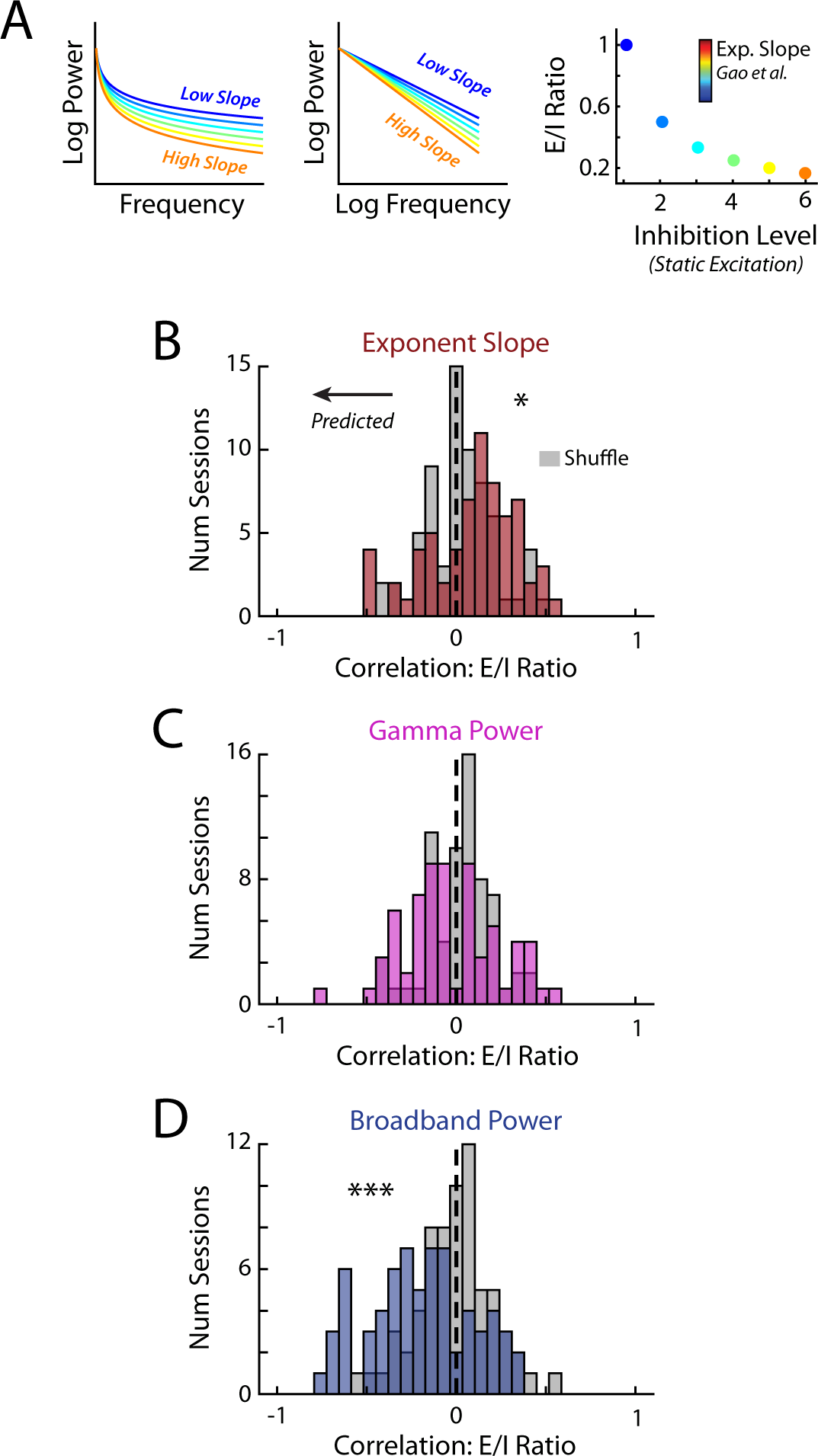
Relationship between LFP and E:I balance. A) A schematic outlining the theoretical relationship between E:I balance and exponent slope as presented in Gao *et. al.*, 2017. Aperiodic spectral components are plotted with varying levels of slope on a linear (left) or log (middle) frequency axis. (right) Varying the level of inhibition (static excitation) alters the E:I balance. As modeled by Gao *et. al.* lower levels of inhibition have shallower exponential slopes. Note that this modeled relationship means that exponential slope is inversely related to E:I balance. (B) Distribution of correlations between the exponent slope and the E:I balance in mPFC as measured by the ratio of the average excitatory synaptic strength against the average inhibitory synaptic strength in each session. CCHs to derive synaptic strength were computed based on the method in Fig 2I from coupled cell pairs identified using the full session data. Note that the significant relationship between Exponent Slope and E:I ratio is *opposite* to that predicted by the Gao *et. al.* model. (C) Distribution of correlations between gamma power and E:I balance. (D) Distribution of correlations between broadband power and E:I balance.

To further understand the relationship between LFP and network communication, we compared our spiking data to other prominent metrics of the LFP. Gamma oscillations (30Hz to 80 Hz) have long been studied as a potential catalyst for relaying and synchronizing information across different brain areas (*37-39*). Gamma power has also been closely associated with interneuron activity (*40-43*), perhaps pointing towards a relationship between LFP gamma power and the balance between excitation and inhibition. Interestingly, even though many models propose a tight link between gamma oscillations and inhibitory dynamics, one side of the E:I balance, we found no significant relationship between the ratio of synaptic excitation and inhibition and the level of gamma power in the LFP (Fig 3C; Mann-Whitney U test vs shuffle; Gamma: z=-1.64, p=0.10, d’=0.24).

Additionally, gross level analyses of overall broadband power within the LFP (2Hz to 150Hz) have been directly linked to neuronal firing rate in patients with epilepsy (*44*). While a relatively crude metric, this past work suggests that broadband power may also provide a lens through which to extract underlying E:I dynamics from emergent LFP signals. Interestingly, we found a strong and significant negative correlation between E:I ratio and overall broadband LFP power over the 1 to 150 Hz frequency range (Fig 3D; Mann-Whitney U test vs shuffle; Broadband: z=-3.78, p<0.001, d’=0.73). Times when LFP exhibited lower spectral power corresponded to times when the E:I balance was skewed towards more excitation and less inhibition. Importantly, this relationship was also apparent when estimating E:I balance based on overall activity of principal cells and interneurons, as opposed to direct synaptic strength (Fig S8C,F). Thus, while our data failed to support past hypotheses describing an inverse relationship between aperiodic slope and E:I ratio, we found strong experimental evidence in support of a link between broadband LFP power and the dynamic balance between excitation and inhibition.

A fundamental understanding of the complex interactions between excitation and inhibition is thought to be critical to the development of effective therapeutics (*9*). Accordingly, numerous models of network functioning and experimental studies have explored the consequences of altering the E:I balance through systematic manipulation of either the excitation or the inhibition component (*7, 45-53*). For example, many models of attractor networks posit that inhibition throughout the network remains stable, serving as a global check, while excitatory drive is more dynamic and acts as the driving factor in variations to the overall E:I balance.

Examining the excitatory and inhibitory coupling in our data independently, we found that inhibitory tone was more tightly linked to ongoing LFP dynamics than excitatory tone (Fig 4A-C; Mann-Whitney U test Excitatory tone vs Inhibitory tone; Exp. Slope: z=1.91, p=0.05, d’=0.09; Gamma: z=-0.03, p=0.97, d’=0.01; Broadband: z=-5.12, p<0.001, d’=0.26). Interestingly, even a simple evaluation of the overall variability (coefficient of variation) in synaptic coupling strength across different magnitudes of LFP components again pointed towards inhibition as a more dynamic process than excitation (Fig 4D-F; Mann-Whitney U test Excite vs Inhibit; Exp. Slope: z=-6.76, p<0.001, d’=0.34; Gamma: z=-6.63, p<0.001, d’=0.35; Broadband: z=-6.97, p<0.001, d’=0.37). This is in direct contrast to most computational models which suggest that changes in excitatory activity drive changes in E:I balance (*46, 48, 49*), but more in line with dynamic changes in a network’s inhibitory tone in response to varying levels of environmental uncertainty (*47*).

**Figure 4:**
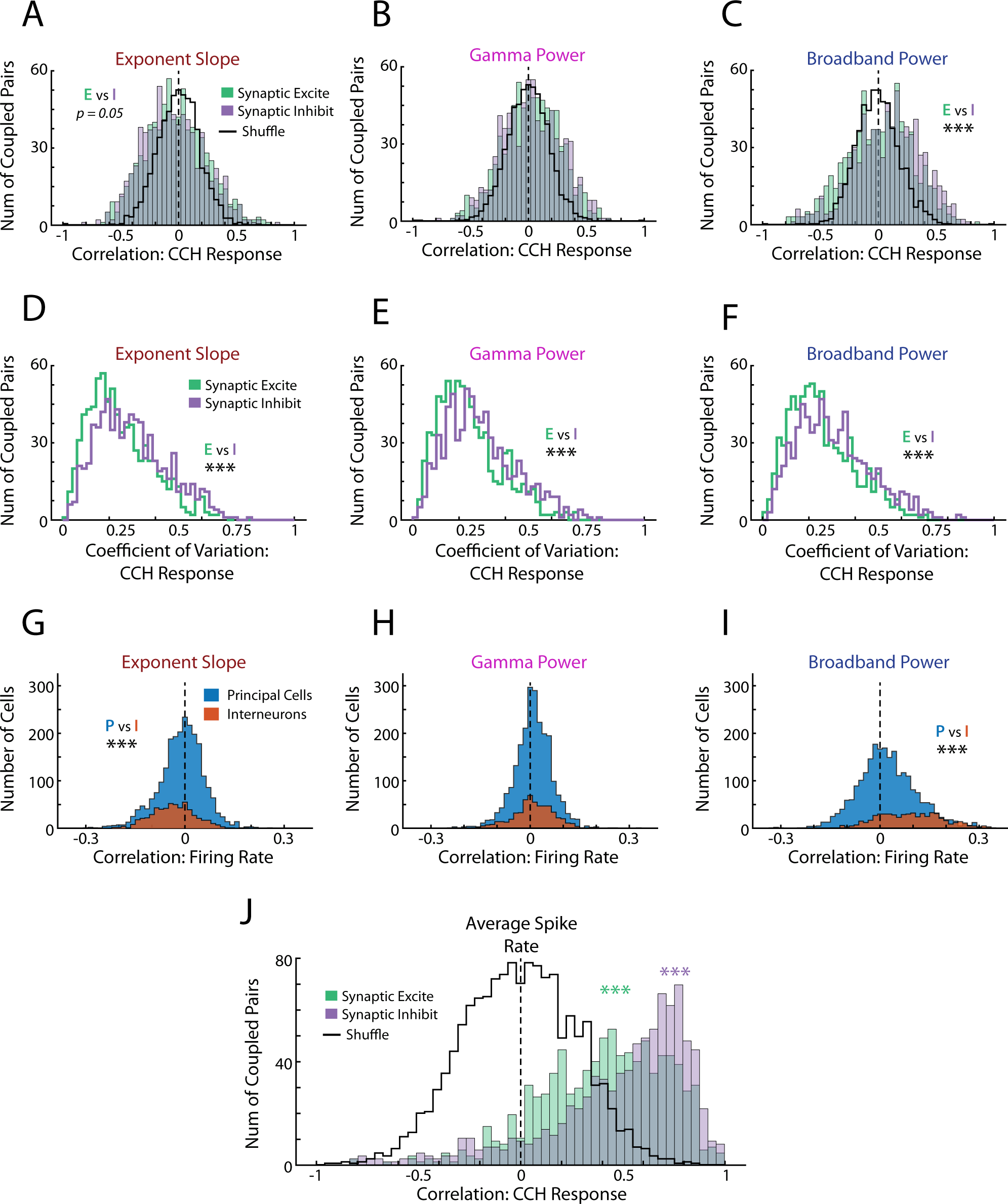
Relating LFP to the independent excitatory and inhibitory components. (A-C) Distribution of correlations between LFP characteristics and strength of synaptic excitation or inhibition. Data as in Fig 3 but using the CCHs from the excitation and inhibition independently. Shuffle depicts the pooled average across the two shuffled data sets. (D-F) Distribution of coefficient of variation of excitatory and inhibitory synaptic response strengths taken across different magnitudes of each of the three LFP characteristics. (G-I) Distribution of correlations between LFP characteristics and average firing rate of principal cells or interneurons. Note that firing rates were computed in each 512ms LFP window yielding a total of ∼8000 LFP-Rate pairs to correlate. J) Distribution of correlations between the average firing rate of a pair of coupled cells and their synaptic coupling strength. CCHs were computed across differing levels of paired firing rates using the method described in Fig 2I.

As a complement to measures of synaptic coupling strength, we also evaluated raw firing rates of mPFC cells directly. Consistent with recordings from human patients (*44*), firing rates were overall more related to broadband LFP power than expected by chance, and skewed strongly towards higher firing rates during periods with greater LFP power (Fig S10, Broadband power vs Firing Rate: d’=0.53 vs shuffle). Further, these relationships between spiking rates and LFP were more pronounced for broadband power as compared to exponent slope or gamma power (ANOVA across metrics: F(2)=349.40, p<0.001; post-hoc HSD Broadband vs Exp. Slop: p<0.001; Broadband vs Gamma: p<0.001), once again suggesting that LFP broadband power may more closely reflect underling network dynamics. Segregating mPFC cells according to their principal cell and interneuron designations revealed that the systematic relationship between firing rate and LFP was more prominent within the interneuron population (Fig 4G-I; Mann-Whitney U test Principal vs Interneuron; Exp. Slope: z=8.41, p<0.001, d’=0.35; Gamma: z=-0.81, p=0.42, d’=0.02; Broadband: z=-14.61, p<0.001, d’=0.74), consistent with our analysis of synaptic coupling strength, and further demonstrating the close relationship between inhibitory signaling and broadband LFP dynamics.

Fascinatingly, we found a strong relationship between the firing rates of each pair of coupled mPFC cells and the strength of the synaptic connection between them (Fig 4J; Mann-Whitney U test vs shuffle; Excite: z=23.02, p<0.001, d’=1.39; Inhibit: z=24.96, p<0.001, d’=1.58), with more prominent synaptic coupling occurring at times where there were overall higher firing rates. Notably, these results are consistent with two independent but complementary theories of network communication. First, studies done primarily in brain slices have highlighted the importance of short-term plasticity (STP) mechanisms in modulating synaptic strengths in responses to firing rate dynamics (*54-57*). Our results are consistent with STP processes *in vivo* in which increases in mPFC firing rates correspond to periods of increased synaptic strength. Second, a wide family of computational models have suggested that dynamic changes in E:I synaptic strengths should parallel changes in network firing rate, increasing or decreasing together, as we see in our experimental results (*21, 58-65*). In these models, this relationship provides stability to neural representations, ensuring that synaptic inputs reliably transfer information even during elevated levels of background activity.

The dynamic balance between excitation and inhibition is centrally important to brain function and dysfunction (*3-12*). Yet, the ability to effectively measure this balance, especially within clinical settings, has remained elusive. Our data present a straightforward solution to this problem, and argue that simple measures of local field potentials can provide insight into the dynamic ratio between excitatory and inhibitory signaling --- broadband spectral power provided the best proxy for the underlying E:I balance. And while we also found that the aperiodic spectrum was related to E:I ratio, their interaction was opposite to previously proposed hypotheses.

Furthermore, our data argue that excitation and inhibition are not equal players in setting the E:I balance, but rather that localized inhibition represents the dominant force and that emergent properties in local field potentials provide a biased lens towards the inner workings of inhibitory dynamics. Thus, we propose that network models should look to inhibitory dynamics, rather than excitatory, as the primary drivers of network changes. Likewise, experimental measures of neural activity and network communication during behavior, such as the naturalistic decision-making studied here, should pay careful mind to the variation in inhibitory signaling as it relates to complex cognition.

Finally, our findings provide critical technical advancements for both fundamental and clinical neuroscience: For fundamental neuroscience, our experiments and analyses revealed that multiple levels of network communication (spiking, synaptic coupling, and emergent LFP) change dynamically in real-world conditions. Large neural ensembles of simultaneously recorded cells in real-world behaviors are becoming easier and easier to achieve (*66-69*) and the techniques developed here are widely applicable across these data sets to probe relationships across these multiple levels in other neural structures.

Clinically, many models suggest that psychiatric conditions emerge as a result of dysfunction in network communication or imbalances between excitatory and inhibitory signaling, particularly in prefrontal cortex (*7-12*). Our findings provide direct experimental evidence for a fundamental link between local field potentials and the balance between excitation and inhibition in the underlying network. As such, the evidence that broadband spectral power is a useful proxy for E:I balance in prefrontal cortex is likely to be particularly useful clinically, even if these relationships have limited generalizability to other brain areas. This work provides a valuable building block in the development of effective diagnostics for clinical practice, arguing that easily recordable and non-invasive signals such as MEG, EEG, and fMRI can provide invaluable access to underlying facets of network communication.

## Methods

### Subjects

All data used in the study were collected from eight adult Fisher-Brown Norway F1 hybrid rats (4M, 4F) on the Restaurant Row task (see Diehl and Redish, 2023 for details). All analyses and findings reported here are novel and were not explored in previous publications. All rats were aged 9-15 months during experiments, were singly housed in a temperature-controlled colony room with a 12-hour light/dark cycle, and were food restricted to at or above their 80% free feeding weight. All procedures were approved by the University of Minnesota Institutional Animal Care and Use Committee (IACUC) and were performed following NIH guidelines.

Data from one rat, R506, were excluded from any analyses involving LFP metrics due to differences in recording hardware and methodologies (see “Recordings”). Although these data (109 of 3017 single units and 448 of 9344 LFP channels) were excluded from the presented results, they were qualitatively similar to all reported findings.

### Restaurant Row task

Rats were trained to perform the economic decision-making task “Restaurant Row” (RRow) as described previously (*16*). Briefly, rats had one hour to collect differently flavored rewards from a set of four serially located food sites (Restaurants). At each site, rats first entered the *Offer Zone* (OZ) where they were cued via the frequency of an auditory tone as to the temporal delay that would wait to receive a reward (1-30 seconds; pseudorandom from a uniform distribution). Rats then had the option to *Skip* the offer and advance to the next reward site, or *Accept* the offer and translocate to the *Wait Zone* (WZ) at which point the delay began to count down, cued by subsequent auditory tones of progressively lower frequency. At the completion of the delay, food was delivered and rats were identified as now in the *Reward Zone* (RZ). After consuming their food reward, rats advanced to the next reward site at their leisure. The area between restaurant sites was designated the *Transition Zone* (TZ) and involved no explicit engagement with the task.

Training on the Restaurant Row task has been described previously. It occurred over the course of 25 days during which progressively longer offer delays were systematically introduced until rats were performing the full version of the task. After implantation of recording hardware rats were retrained using an expeditated sequence over the course of 10 days. All recordings were collected after rats were well trained on the task and behavioral responding was stable.

### Surgery

Implantation of silicon probes targeting the medial prefrontal cortex (Acc, dPL, vPL, IL) has been described previously (*16*). Briefly, rats were anesthetized with isoflurane (2% in O_2_) and implanted with either one or two independently movable silicon probes targeted along the medial wall of the medial prefrontal cortex (mPFC). Probes targeted either the dorsal or ventral half of the medial wall. Rats receiving dual implants had one probe targeted to each hemisphere and depth (e.g. Dorsal Right and Ventral Left). All targeting of hemisphere and probe depth was counterbalanced across rats, accounting for sex, but did not consider any behavioral metrics. During surgery rats received carprofen (5mg/kg, SC), dual-cillin (0.2ml, IM), and saline (3ml, SC) for analgesia, prophylaxis, and hydration respectively and at the completion of surgery were given an injection of Baytril (10mg/kg, SC), additional saline (20ml/kg, SC), and oral administration of Children’s Tylenol (0.8ml). All drugs were provided by UMN Research Animal Resources.

### Recordings

Recordings were performed over 14 days (7 days for R506) and have been reported previously (*16*). All recordings included the 1-hr behavioral session, as well as approximately 5 min pre/post periods of quiet rest away from the task. All analyses considered this full set of recording time, about 70 minutes in total.

Recordings from R506 were collected using a Neuralynx 96-channel Digital Lynx SX recording system and a custom built adaptor to join the 64 channel Omnetics based silicon probe to a pair of 32-channel Neuralynx headstage connectors. Analog signal was passed from the rat, through the adaptor, to the pair of 32 channel headstages where it was preamplified 10,000x. Analog signal continued through a commutator to the recording system where it was digitized at 32kHz. For the remaining rats, recordings were collected using a 256-channel Intan RHD recording system connected to 64-channel Intan RHD headstages. For these rats, data were digitized at the headstages at a sampling rate of 30kHz and were then passed as multiplexed digital signals through a commutator to the recording system. Because of the technical differences between these recording setups (sampling rate, analog vs digital signal transfer, preamplification, custom adaptor, recording system), measures of local field potentials were quantitatively different for rat R506 as compared to the other subjects (Fig S5H-J). While all relational metrics and qualitative effects were comparable between R506 and other subjects, we excluded rat R506 from any formal analysis involving LFP due to these differences in recording methodology.

Single unit identification was performed after 600Hz high-pass filtering of the raw data using Kilosort v2.0 (https://github.com/MouseLand/Kilosort) (*70*), with manual post clustering curation performed using Phy v2.0 (https://github.com/cortex-lab/phy). Local field potentials (LFP) were extracted by down sampling raw recordings using the Matlab function ‘decimate’ to a sampling rate of 2000Hz and low-pass filtering at 800Hz.

After completion of all recordings, rats were transcardially perfused and histological verification confirmed all recording sites, as previously reported (*16*). This allowed us to precisely localize all single units and LFP channels in a standardized 3-dimensional space. Only recordings from the medial prefrontal cortex were analyzed subsequently: 3017 single units (109 from R506) and 9344 LFP channels (448 from R506).

## Data Analysis

### Behavior

Extensive analysis of behavioral responding from these subjects has been performed previously (*16*). Behavior was largely consistent with other implementations of the Restaurant Row task (*71-73*) and responses were consistent with the use of task information and well-founded economic strategies. All recording times were categorized according to task period as being in the Offer Zone (OZ), Wait Zone (WZ), Reward Zone (RZ), or Transition Zone (TZ), or in the Pre or Post task period.

### Cross-Correlation Histograms (CCHs)

For all pairs of simultaneously recorded cells we computed cross-correlation histograms (CCHs) in 0.5ms time bins over a ±150ms window around the reference spike times of a Cell A (*17, 18*). This yielded the total number of spikes from the receiver Cell B at times relative to the spiking of a driver Cell A (Cell A → Cell B). Note that for interacting cell pairs, this will yield temporally precise deviations, either increase or decreases, in the spiking rate of Cell B. To identify these significant deviations we first computed the expected CCH relationship in the event of no short timescale relationships by convolving the CCH with a 25ms gaussian kernel (*74*). This method is largely equivalent to performing a temporal jittering procedure on each spike (12.5ms jitter), and serves to destroy short time scale relationships between the cell pairs (<jitter window) while maintaining long time scale relationships (*74, 75*). Confidence bounds around the convolved CCH were then computed as the bounds of a Poisson distribution, taken at the α = 0.001 level. Significant CCH responses were identified as bouts of at least 3 consecutive time bins (1.5ms) that exceeded the confidence bounds. Significant responses in the same direction that occurred less than 3 bins apart (within 1.5 ms) were joined together as a single event. Positive and negative responses were computed independently for each CCH and each event was taken as the net deflection beyond the respective confidence bound. For each identified CCH deflection, the response time was taken as the center of mass of the deflection and response magnitude for the base CCH was the sum of the net response normalized by dividing by the median spike count across the full time window. Based on the empirical distribution of response times (Fig S3A) and past work (*18, 57, 76, 77*), we did not consider any coupled pairs with a response time beyond 6ms.

Because CCHs are bidirectional and computed for each cell pair in each direction, a relationship from Cell A to Cell B in the positive time direction would also appear mirrored as a relationship from Cell B to Cell A in the *negative* time direction. Thus, for subsequent analyses we restricted our set of identified CCH responses to only those that described a *forward* (positive time) relationship from Cell A to Cell B. Additionally, because true positive Cell A to Cell B responses should appear in mirror when comparing Cell B to Cell A, we restricted our analysis to only forward responses in which our CCH analysis also detected a reverse response with a comparable time offset and magnitude profile (within 2ms time and 50% response magnitude). All subsequent analyses included only those cell pairs for which we detected a significant, forward coupling relationship based on the full recording data (3142 coupled pairs).

For each identified forward relationship, we classified the response direction as either excitation or inhibition according to increases or decreases in spiking rates relative to the convolution derived baseline. Amongst excitatory couplings, response times followed a clear bimodal distribution with a local minimum between them at approximately 1ms (Fig S3B). Thus, we segregated identified coupling relationships into those that were “Co-Spiking” vs those that were putatively monosynaptic according to whether the response time was less than or greater than 1ms. Note that while inhibitory connections could also be identified as “Co-Spiking” in that there was a significant decrease in spiking from Cell B concurrent with spikes from Cell A, this represented a very small proportion of the data and was not explored further. Thus, our CCH analysis yielded three prominent responses categories: Synaptic Excitation (positive deflection between 1ms and 6ms), Synaptic Inhibition (negative deflection between 1ms and 6ms), and Excitatory Co-Spiking (positive deflection at <1ms). Note that at no point were these categories considered when identifying CCH responses, nor were there any limitations baring a given cell pair from being identified as a member of multiple categories.

For analyses in which we computed CCHs over specific task conditions (behavioral phases or LFP quantifications) we examined only those cell pairs with an identified coupling based on the full recording data. When restricting CCH time windows, either for computing during behavioral epochs or during chunks of aggregate time based on LFP characteristics or firing rates, only the spike times of the input cell were restricted to the period of interest. Spikes from the recipient cell were considered regardless of restriction conditions and needed only to fall within the window for the CCH analysis (±150ms). After restriction, CCHs were z-scored to normalize for potential differences in baseline firing rates, and the response magnitude was taken as the average z-scored CCH within a window of interest. Note, no comparison to jittered confidence bounds was used for these restricted CCHs. For synaptically coupled pairs, coupling strength was taken as the z-scored CCH mean between 1ms and 6ms, and for co-spiking coupling strength was taken as the CCH mean between −1ms and +1ms.

### Cell class designations

Classification of single units into putative “Principal Cells” and “Interneurons” was made according to average firing rate during behavior and the time between the peak and valley of a cell’s average waveform; its spike width (*16*).

### Characterization of multi-step network relationships

To characterize multi-step synaptic interactions in mPFC we examined the putative synaptic inputs into cells for which our CCH analysis had identified that the candidate cell provided significant synaptic drive. For each identified input cell we examined all other cell pairs to determine if any other recorded cells provided synaptic input to the candidate cell of interest. Of those candidates in which we identified at least one other mPFC cell that provided synaptic drive, we further identified the proportion of candidate cells that received at least one excitatory synaptic connection and those that received at least one inhibitory synaptic connection. Note that all candidate cells either *received* or *did not* receive synaptic input from other mPFC cells. Of those cells that received input, this could come from *both* excitatory and inhibitory connections.

### LFP quantifications

Spectrograms of all LFP channels were computed using a Short-Time Fourier Transform (“spectrogram” in Matlab) in 512ms chunks using a 1024 sample hamming window. Windows started from the first recording sample with no overlap between windows and were computed irrespective of any ongoing behavioral considerations. Each spectrogram was computed between 1Hz and 150Hz at 0.5Hz intervals, the real component was extracted, and the log10 power in each sample was used for all subsequent analyses. For each spectrogram we quantified 3 parameters: exponential slope, gamma power, and broadband power. First, each spectrogram was converted into log-log space such that the underlying exponential aperiodic component was linear over frequencies. Next, we computed a robust linear fit of the log-log spectrum using the Matlab function “robustfit”. Note that this linear fit in log-log space is mathematically equivalent to an exponential fit in linear-frequency space, but markedly more computationally efficient. From this linear fit, we took the exponential slope as the inverse of the fit slope parameter (equivalent to the exponential decay rate in linear-frequency space), and the broadband power as the average power of the fit spectrum. Gamma power was taken as the mean power in the gamma range (35Hz – 95 Hz, see below) relative to the fit spectrum. Note that this method ignores the influence of sustained oscillations on the spectral power, though as can be observed in the example, spectrum computed on these limited time windows were noisy around the aperiodic spectral power and generally lacked obvious oscillatory components (Fig 2B). Thus, we argue that our method of a robust linear fit provides a viable estimate of underling LFP spectral components on these short timescales (Fig S5). In support of this conclusion, analysis of each 512 ms spectrogram using the FOOOF package to explicitly decompose spectrogram into aperiodic and oscillatory components yielded highly comparable results on an example session (Fig S5E-G). Furthermore, we also computed the average power spectral density over the full recording time and repeated both our simple robust linear fit based calculations and FOOOF based aperiodic extraction (Fig S5A-D). Both of these methods produced quantifications of our relevant LFP components over the full recording data that were comparable to distributions across 512 ms spectral windows.

Fitting of all spectrograms with the FOOOF analysis method (*14*) used a Matlab iteration of the original approach written by Luc Wilson (*78*). Data were fit over the full computed spectrum from 1Hz to 150Hz. The aperiodic component was fit using the “fixed” method with no knee, a maximum of 8 oscillatory peaks were allowed, the minimum peak height was 2std above the aperiodic, and gaussian peak standard deviations were limited to between 2 and 20 Hz.

To identify a suitable frequency range for gamma oscillations in our mPFC data we used the FOOOF decomposition method to extract oscillatory power and identified peaks across all LFP channels. From these data we identified the gamma oscillation frequency in our mPFC data to range from 35Hz to 95Hz. Our data had no clear evidence of distinct high and low frequency components of the gamma oscillation, thus we use a singular range for gamma power.

### Generation of chunked CCHs

In order to compute CCHs related to ongoing LFP characteristics we performed an aggregate chunking procedure that grouped periods of spiking data that occurred during similar bouts of LFP activity. First, LFP spectrograms were computed at 512ms bins and quantified as described above. Second, the distribution of quantifications (e.g. the exponent slope) was collected across the full recording (∼8000 samples over ∼70min recording) and time bins were discretized into 20 equal groups according to their percentile across the full distribution. Note that while LFP channels were very highly correlated (Fig S6), this discretization differed mildly according to the specific LFP channel that was used. As described below, this process was repeated in triplicate with randomly selected channels to minimize the impact of spurious results. Third, spiking data from time bins in each group/percentile range were aggregated to yield a total of ∼3min of data. Fourth, CCHs were computed for each set of aggregated spiking data. This yielded a total of 20 CCH results for each pair of coupled cells, with each CCH corresponding to differing levels of exponent slope (or the LFP characteristic of interest). Fifth, each CCH was normalized by z-scoring to account for potential differences in baseline firing rates. Sixth, the response magnitude was calculated for each aggregated CCH by taking the mean z-scored values across a window of interest. For synaptically coupled pairs, this was between 1ms and 6ms, and for co-spiking pairs, this was between ±1ms.

For comparison of CCH responses to each LFP characteristic this aggregate procedure was used, with the discretization of the time bins based on the values of interest (exponent slope, gamma power, or broadband power). For relating CCHs to firing rates, we first computed the average firing rate for each cell in a given pair in the same time bin windows that were used for LFP calculations. For each time bin we then took the mean firing rate between the pair of cells as the overall level of spiking activity in that time window for that cell pair. Discretization of time bins was performed on the distribution of pair averaged firing rates, and CCH calculations were then performed as for LFP characteristics.

For calculation of variation in excitation or inhibition we computed the coefficient of variation of synaptic coupling strength for each coupled pair. CCHs were computed from aggregate time chunks as described above yielding 20 coupling strengths for each cell pair. Coefficient of variation was then computed by dividing the standard deviation across these 20 values by their mean.

### Relation of CCHs/firing rates to LFP

To relate CCH coupling strength to LFP characteristics we computed sets of 20 chunked CCHs as described above. For each coupling pair the set of chunked CCHs were computed in triplicate for three randomly selected LFP channels from the same Si probe that recorded the unit data. For each given session the same sets of LFP channels were used to evaluate all coupled cell pairs. After computing the coupling strength via CCHs, this vector of 20 strength values was correlated to the mean LFP characteristic in each aggregate time chunk. We then took the median correlation between CCH coupling and LFP characteristic across the three independent LFP channels as the quantitative relationship between the coupling strength of the given cell pair and LFP dynamics. Comparison of the relationship between CCHs and cell firing rates followed the same process with chunked CCHs aggregated based on discretizing firing rates across time bins, as described above.

When comparing cell firing rates to LFP, we computed the average firing rate of the given cell in each 512ms time window used for computing LFP spectrograms. This vector of firing rates (∼8000 samples) was then correlated to the LFP quantification across a set of three randomly selected LFP channels. As above, candidate LFPs were recorded on the same Si probe as the cell of interest, and the final quantification was taken as the median correlation across the triplicate repeat.

### Measures of Excitation:Inhibition ratio

To relate LFP metrics to the balance between Excitation and Inhibition we computed three alternative measures of the Excitation:Inhibition Ratio:

1. *Ratio between synaptic excitation and synaptic inhibition*. Coupling strength was computed for all identified synaptically coupled pairs and the mean excitatory response was divided by the mean inhibitory response within each session to yield E:I ratio. This process was performed independently for sets of chunked CCHs (see above), yielding a set of 20 E:I ratios to compare against LFP metrics.
2. *Ratio between principal cells (excitatory) and interneurons (inhibitory)*. Firing rates of all cells were computed in 512ms bins corresponding to quantification of LFP data. For each time bin, excitatory input was taken as the mean firing rate across all identified principal cells and inhibitory input as the mean firing rate across all identified interneurons. E:I ratio was computed in each bin by dividing excitatory activity by inhibitory activity.
3. *Ratio between cells with identified synaptic excitation and those with identified synaptic inhibition*. Firing rates for all recorded cells were again computed in 512ms bins corresponding to LFP. From the full set of cells, excitation was then taken as the mean firing rate across all cells in which CCH analysis identified at least one excitatory synaptic connection and inhibition was the mean firing rate across all cells with at least one identified inhibitory synaptic connection. E:I ratio was then taken as average excitation divided by average inhibition in each 512ms bin.

For all metrics, the computed E:I ratio was then correlated against different characteristics of the LFP, either across 20 chunked CCHs (measure 1) or approximately 8000 time bins of 512ms (measures 2 & 3). Only recording sessions that contained at least one excitatory and one inhibitory candidate (cell pair or individual cell) were included in analyses. For each session, 3 LFP channels were selected at random from each recording probe and the E:I ratio was correlated to each of the three. The median correlation was taken for each Si probe to reduce the impact of spurious results. For rats with 2 recording probes, each probe was treated separately, correlation between LFP and E:I ratio and was computed for each, and the median across the two probes was taken for the given session as the relationship between LFP and E:I ratio.

### Statistics

Comparison of LFP characteristics and CCH coupling strengths across different task periods was performed with a one-way ANOVA. Comparisons of proportion of cells were done with a Chi-squared test for equal proportions. Comparison of the correlation between LFP characteristics and E:I balance or firing rate was done with a Mann-Whitney U test against a null distribution generated by shuffling cell inter-spike intervals (ISIs). To generate this null shuffle, the ISIs of each mPFC cell was randomly permuted to dissociate spiking from behavior, LFP, and the spiking of other cells. ISIs of each cell were permuted independently. Shuffles were performed one time for each cell to generate a single distribution of shuffled analysis results against which real data were compared. Comparisons between synaptic excitation and synaptic inhibition or between principal cells and interneurons were done with a Mann-Whitey U test. Correlations used a Person’s correlation, and all statistics were evaluated at an α = 0.05 level. Effect sizes (d’) for two sample comparisons are reported in line with statistical results. These were calculated in the standard manner as the absolute difference between the group means divided by the pooled standard deviation across the two groups. All statistical comparisons were performed on data grouped across all rats, though in all cases qualitative results held for each rat individually.

For all display and statistical purposes the number of samples was as follows: For comparisons of synaptic couplings sample number was the number of identified coupled pairs. For cell firing rates, sample number was the total number of mPFC cells. For E:I balances, sample number was the total number of recording session with at least one excitatory and one inhibitory component.

## Data and Code availability

All analyses occurred in Matlab v. 2017b. Upon publication, all datasets and analysis code used in the present study will be uploaded to an appropriate public repository.

## Acknowledgments

We thank K. Seeland, C. Boldt, A. Sheehan, and C. Bogner for technical assistance as well as members of the Redish lab for useful discussion. This work was supported by NIH grants R01-MH112688, P50-MH119569, and T32-MH115886.

## Author Contributions

G.W.D and A.D.R designed the study. G.W.D. collected and analyzed the data. G.W.D and A.D.R wrote the manuscript.

---

**Figure S1:**
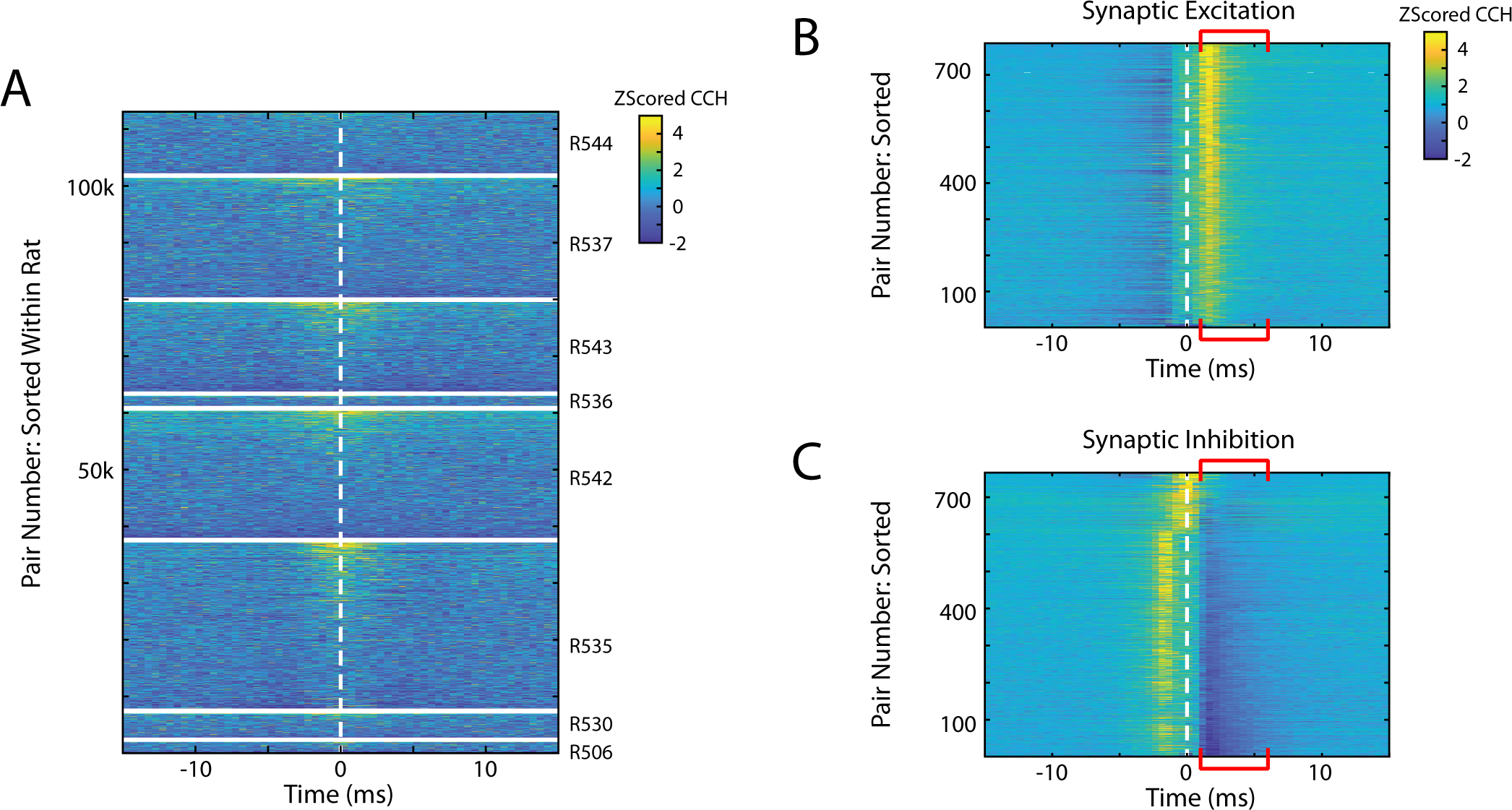
Full set of CCH couplings. A) Z-Scored CCHs for all 107418 pairs of simultaneously recorded mPFC cells. Cells are grouped by rat and sorted within each rat according to the mean value between ±5 ms. Note that gaps have been added between each rat. (B) CCHs of all cell pairs with an identified excitatory synaptic connection. Pairs are sorted according to their coupling strength. Red brackets denote the time window for considering synaptic interactions. (C) CCHs of all cell pairs with an identified synaptic inhibitory connection.

**Figure S2:**
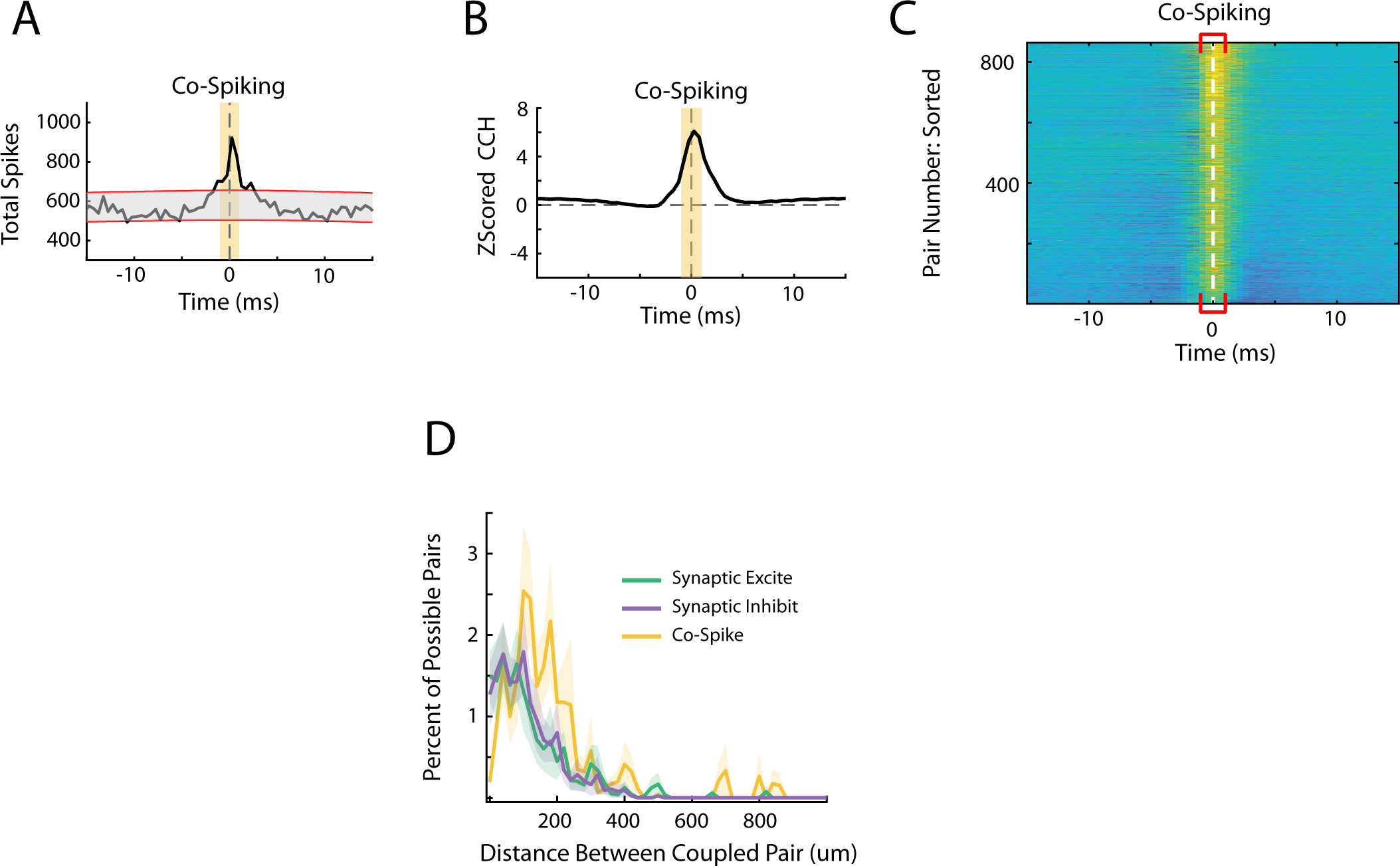
Co-Spiking in mPFC. A) An example cell pair that exhibited zero-lag co-spiking (*R535-2019-07-21 SiC1_07 ◊ SiC1_14*). Yellow shading represents the time window for identifying co-spiking. B) The average CCH for all co-spiking cell pairs. C) CCHs of all identified co-spiking pairs. D) The proportion of identified coupled pairs and the anatomical distance between them. Synaptic data are replotted from Fig 1H with co-spiking pairs now included. Pearson’s correlation for co-spiking proportion vs distance: r=-0.79, p<0.001.

**Figure S3:**
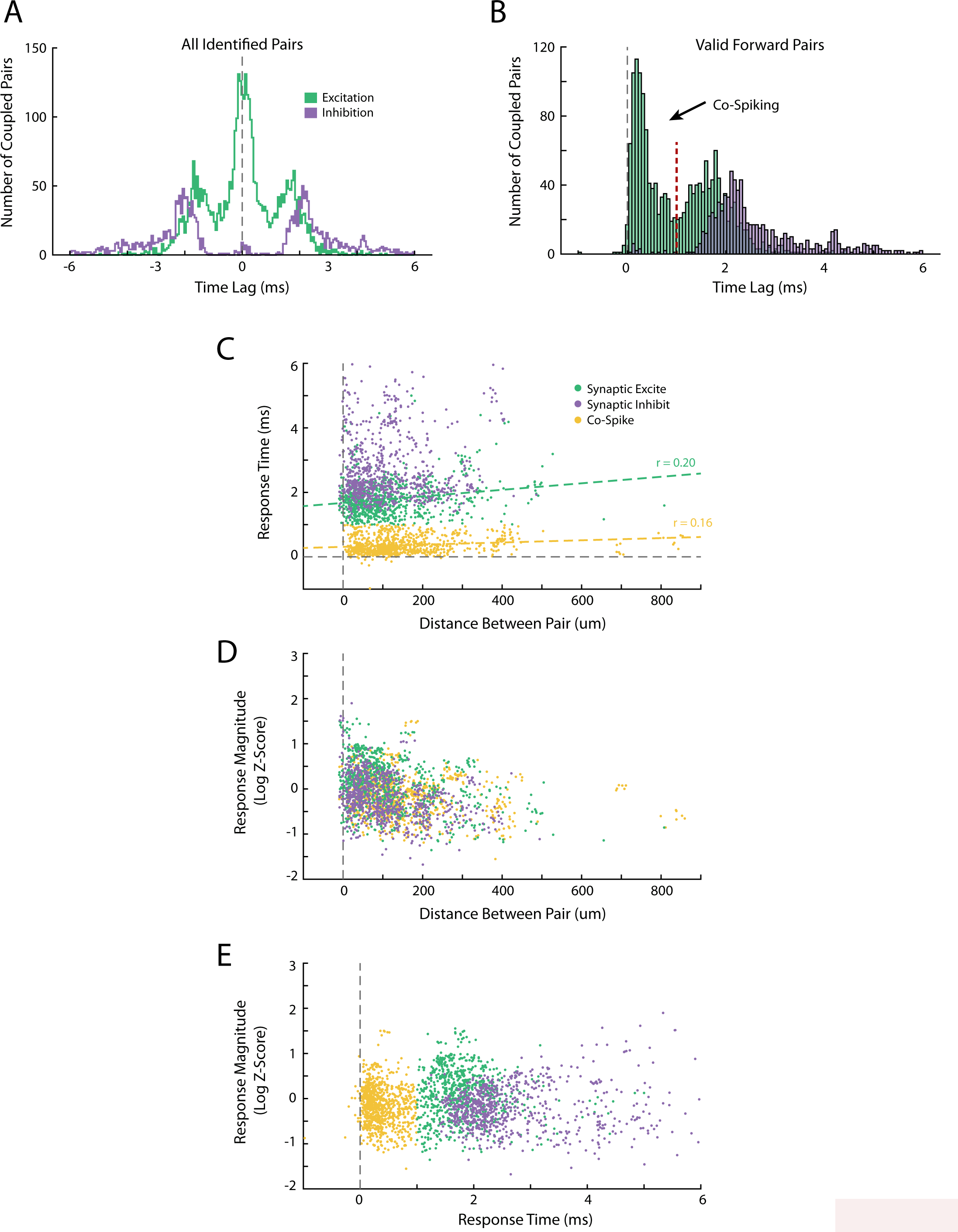
Relationships between CCH properties. A) Distribution of response times for all significant CCH deflections. Note that this plots both forward (positive time) and reverse (negative time) direction interactions between cell pairs. B) Distribution of response times for only those paired couplings that were identified as valid forward couplings (see methods). A clear local minimum was observable for excitatory couplings at approximately 1ms lag. Subsequent analyses used a 1ms boundary to divide co-spiking from synaptically coupled cell pairs. C) The relationship between anatomical distance between a coupled cell pair and the response lag of its coupling relationship. There was a significant positive correlation for excitatory synaptic couplings and co-spiking pairs (Pearson’s correlation; Excite: r=0.20, p<0.001; Inhibit: r=0.06, p=0.09; Co-Spike: r=0.16, p<0.001). D) Relationship between the strength of the coupling response and the distance between the cell pair. E) Relationship between the strength of the coupling response and the temporal lag of the response.

**Figure S4:**
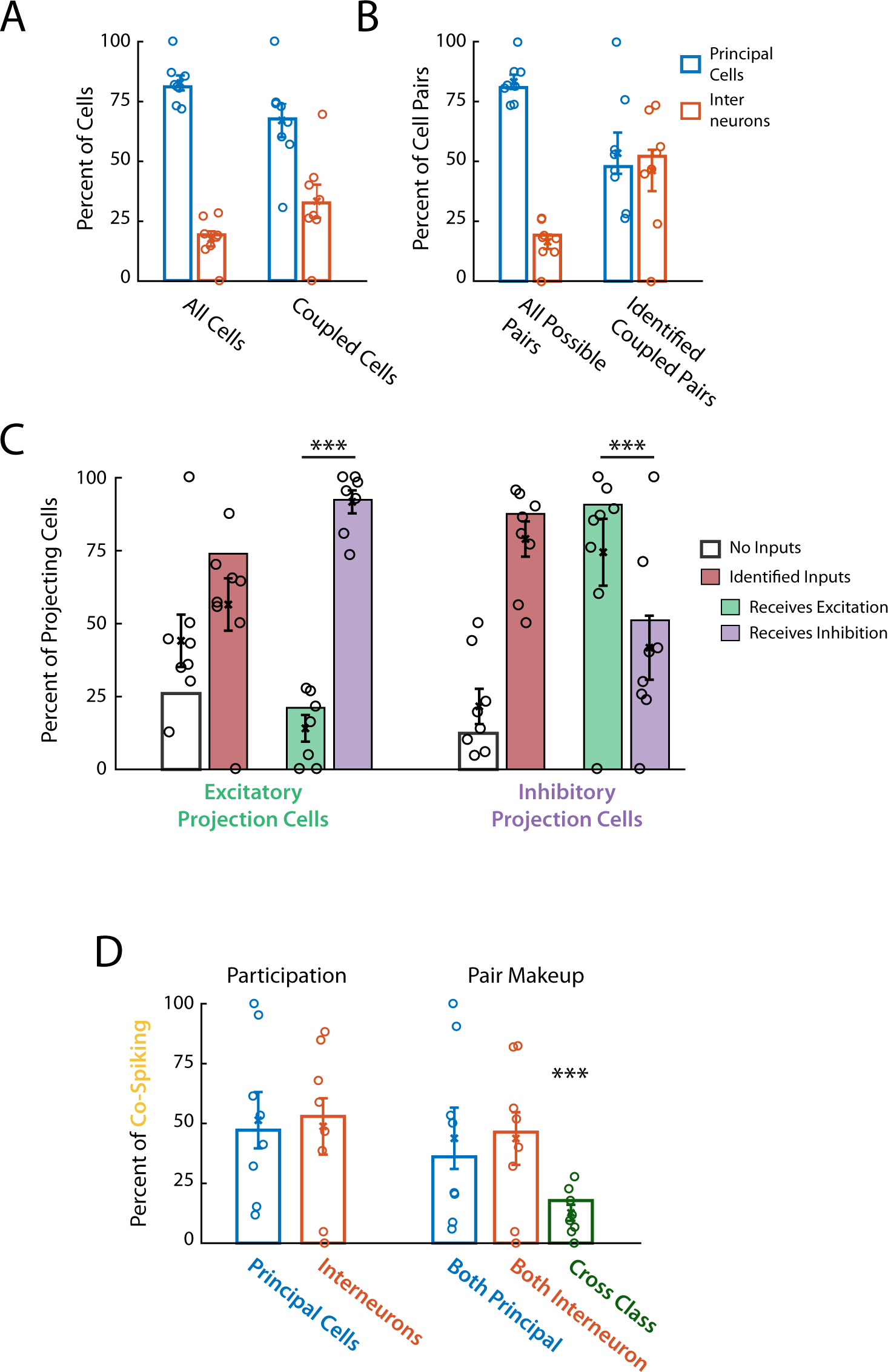
Cell couplings for principal cells and interneurons. A) Proportion of principal cells and interneurons across the full data set or across those cells with at least one identified coupling relationship. Note that interneurons were over-represented in coupling relationships. B) Proportion of all possible cell pairs or those that were identified as significantly coupled which contained a principal cell or interneuron. Again, interneurons were over-represented amongst coupled pairs. C) Breakdown of the inputs onto mPFC cells which were identified as providing excitatory or inhibitory synaptic drive. For each cell identified as delivering at least one excitatory synaptic drive (left bars), we evaluated all other cell pairs to identify if the candidate cell received synaptic input from any other simultaneously recorded cells (open and red bars). For those candidate cells which received at least one synaptic input (red bar), we identified if the candidate cell received any synaptic excitation or synaptic inhibition (green and purple bars). Note that candidate cells could receive both excitation *and* inhibition. This process was also performed for those cells with at least one instance of delivering synaptic inhibition (right bars). This analysis revealed that most cells providing a synaptic drive also received synaptic input from other mPFC cells. Further, cells providing synaptic excitation were more likely to receive inhibition and those providing synaptic inhibition were more likely to receive excitation (Chi^2^ test; Excite: Χ^2^=602.9, p<0.001; Inhibit: Χ^2^=255.7, p<0.001). D) Breakdown of the participation of principal cells and interneurons in co-spiking cell pairs. Co-Spiking pairs were more likely to be both principal cells or both interneurons as opposed to one of each class (Chi^2^ test; vs. both Principal: Χ^2^=73.6, p<0.001; vs. both Interneuron: Χ^2^=162.2, p<0.001). For all plots, data are shown as in Fig 1I. Proportions across the full data set are shown as bars and statistics were calculated on proportions across the full data set. Proportions from each rat are shown as circles and error bars denote the mean ± SEM proportion across rats.

**Figure S5:**
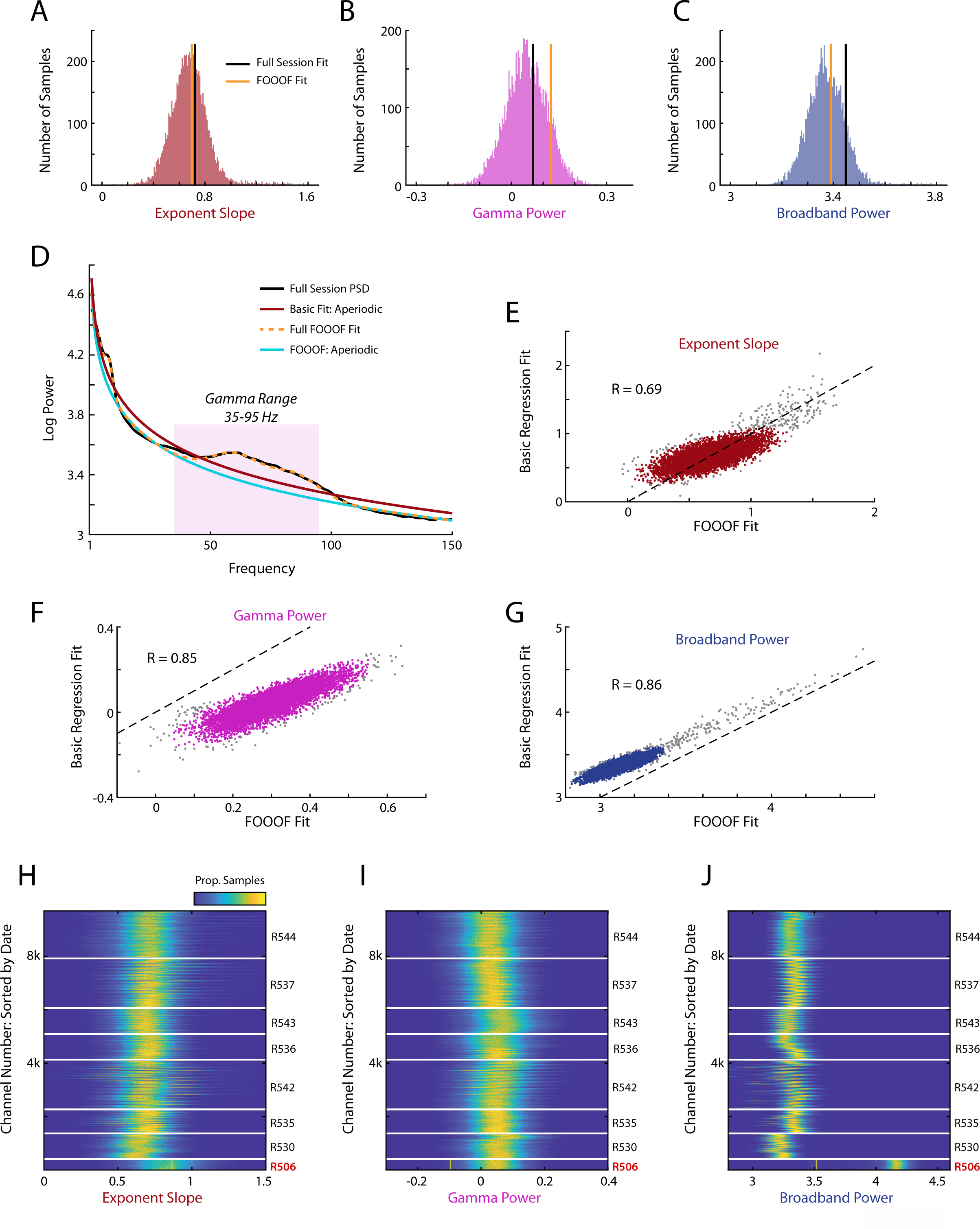
Quantification of LFP components. (A-C) Distribution of exponent slope (A), gamma power (B), and broadband power (C) for an example session (*R542-2019-07-21 SiC1_*14) with spectra computed in non-overlapping 512ms windows. D) Spectral power across the full session was computed by averaging all 8314 windows of 512ms, comparing quantification of the three LFP components of interest using both a basic regression fit and the FOOOF spectral decomposition method. Full session quantifications are plotted in (A-C) as black and orange lines. (E-G) For each 512ms window of the example session we computed each LFP component of interest using a basic regression fit and the FOOOF method. For all three components the two methods were highly correlated (Pearson’s correlation; Exp. Slope: r=0.69, p<0.001; Gamma: r=0.85, p<0.001; Broadband: r=0.86, p<0.001). Outliers were identified according to 2D Mahalanobis distance at α = 0.01 (grey points) and are excluded from these correlations. (H-J) Distribution of all three LFP components across every recording channel. Recording channels are sorted chronologically within each rat. Note that data from R506 were quantitatively different from the other subjects and are thus excluded from subsequent analysis (see methods).

**Figure S6:**
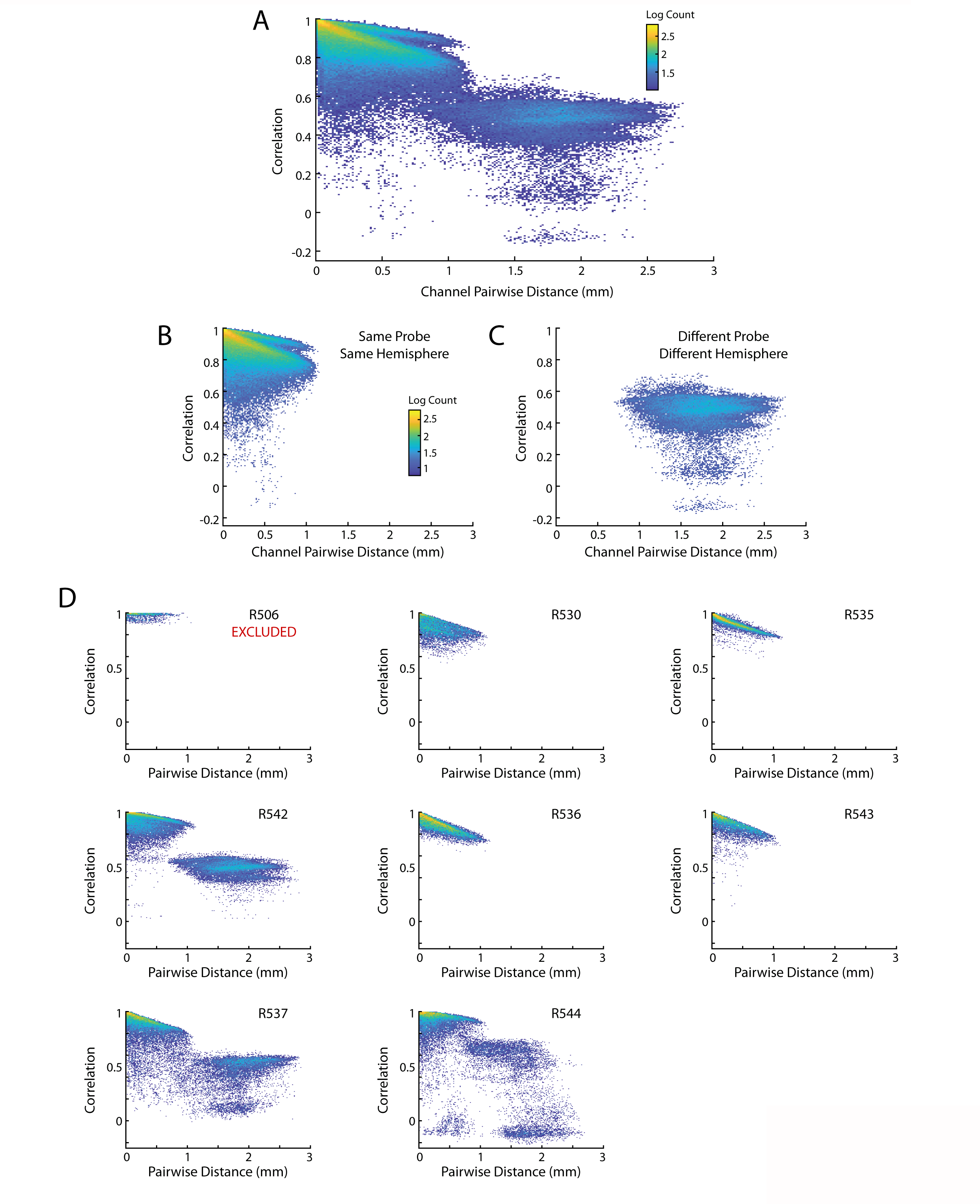
Correlation of LFP as a function of distance. A) Pairwise correlation of simultaneously recorded LFP channels as a function of anatomical distance between the recording sites. Data are binned and presented as a heat map, colored by the log10 count. Bins with fewer than 10 data points are omitted from the plot. (B,C) Data from (A) is replotted but now separated according to if the pair of LFP channels were recorded from the same probe/hemisphere or a different probe/hemisphere. Within a given probe/hemisphere LFP recordings were very highly correlated (typically r>0.8). Yet, correlations decreased with increasing distance between the recording sites. LFPs taken from different probes/hemispheres were less correlated and also did not exhibit the same strong dependence on anatomical distance. Pairwise comparison of LFP quantifications (exponent slope, gamma power, broadband power) yielded comparable results (data not shown). D) Pairwise correlation of LFP plotted for each rat independently.

**Figure S7:**
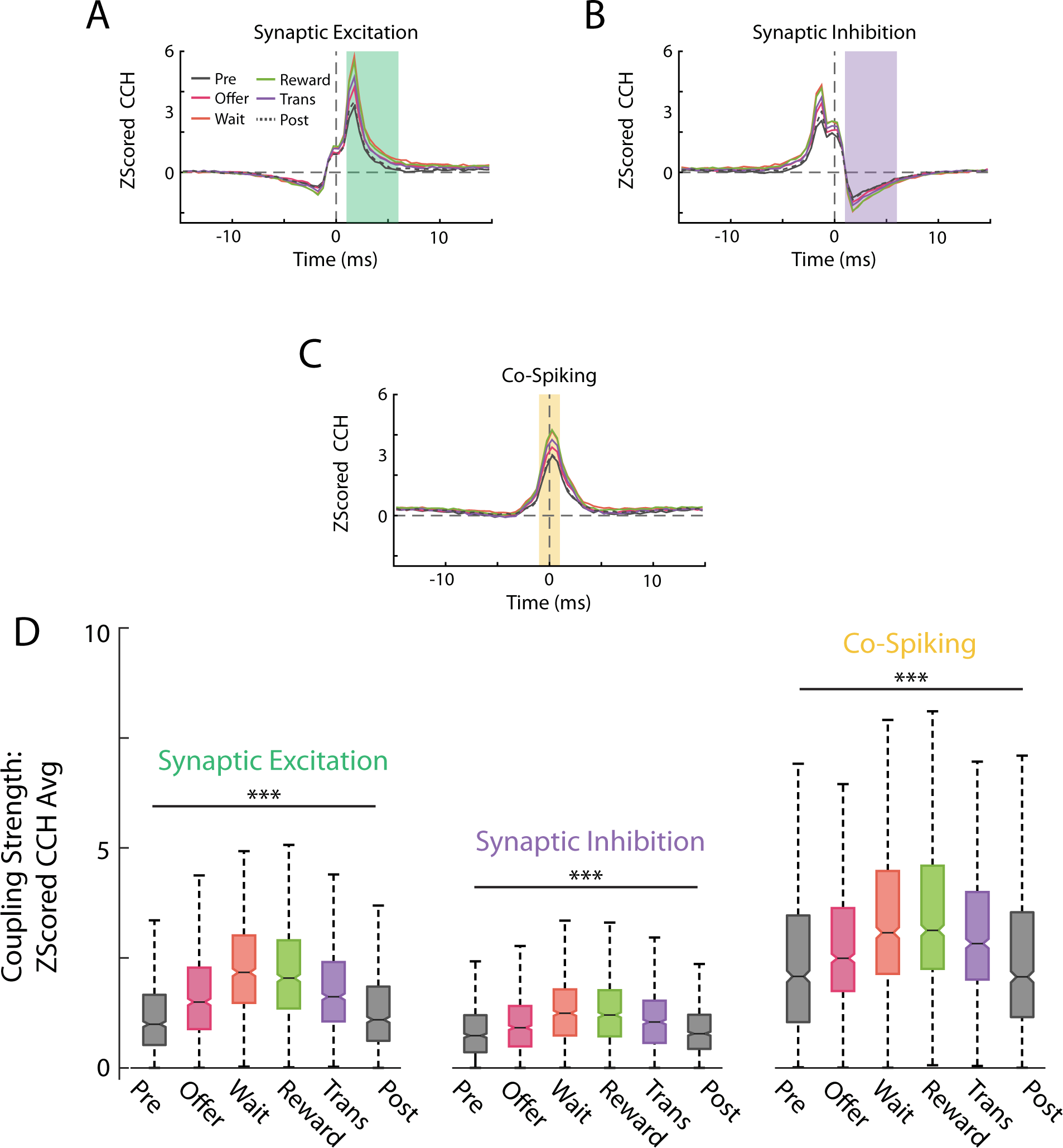
Cell coupling across behavior. (A-C) Average z-scored CCHs for synaptically excited (A), inhibited (B), or co-spiking (C) cell pairs across different phases of the Restaurant Row task (see Methods). D) Distribution of coupling strengths across behavior. Co-Spiking strength differed across the behavioral task (ANOVA across behavior; F(5)=57.3, p<0.001). Data from excitatory and inhibitory synaptically coupled pairs are replotted from Fig 2G,H for comparison

**Figure S8:**
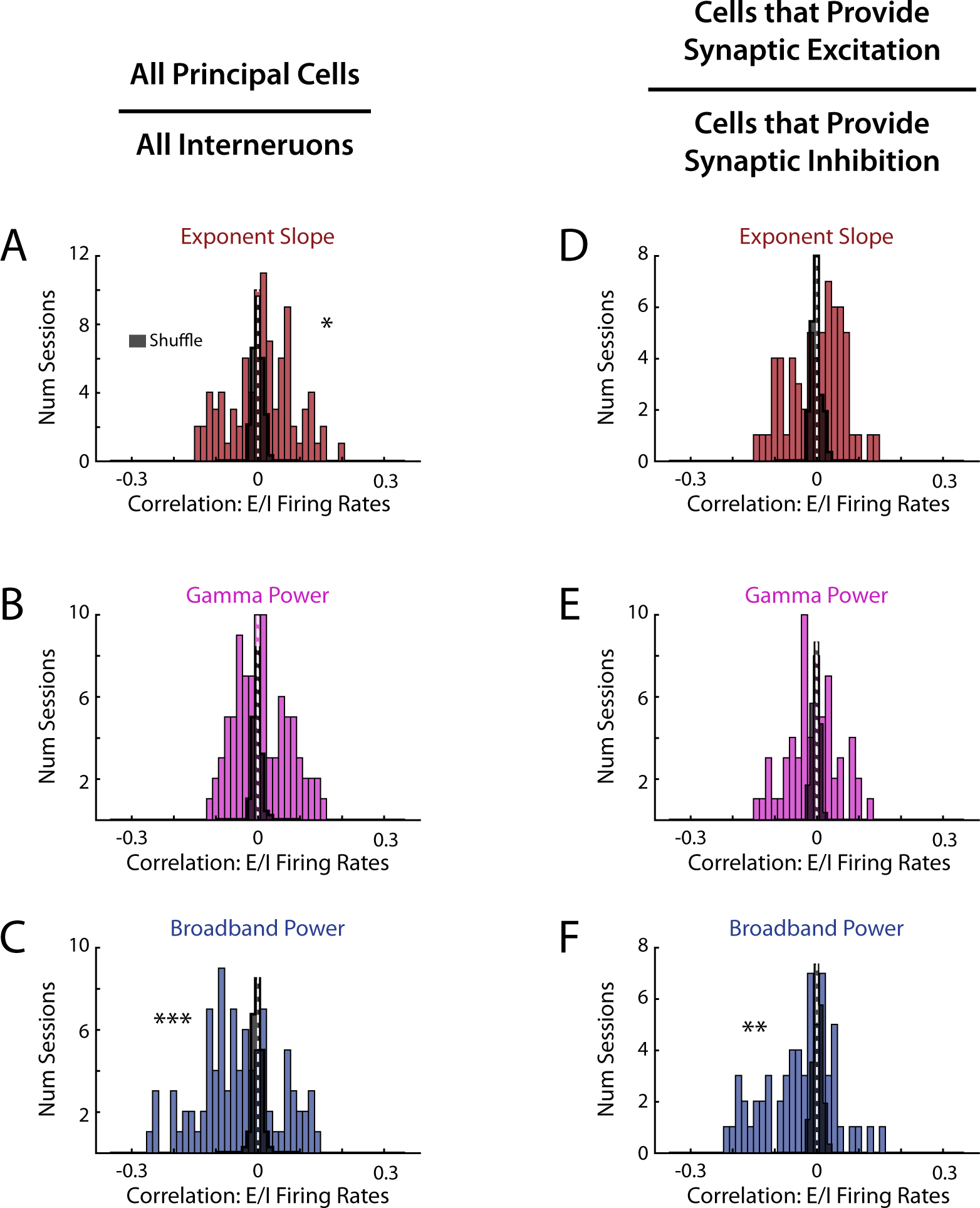
Alternative methods for relating E:I balance to LFP. (A-C) Comparisons taking E:I balance as the mean firing rate of all principal cells divided by the mean firing rate of all interneurons. A) Distribution of correlations with exponent slope (z=2.02, p=0.04, d’=0.19). B) Distribution of correlations with gamma power (z=0.91, p=0.36, d’=0.24). C) Distribution of correlations with broadband power (z=-4.28, p<0.001, d’=0.64). (D-E) Comparison taking E:I balance as the ratio between firing rate of cells with an identified excitatory synaptic connection and those with an identified inhibitory synaptic connection based on CCH analyses. D) Distribution of correlations with exponent slope (z=1.00, p=0.32, d’=0.02). E) Distribution of correlations with gamma power (z=-1.19, p=0.23, d’=0.17). F) Distribution of correlations with broadband power (z=-3.03, p=0.002, d’=0.70). All stats are Mann-Whitney U test vs shuffle.

**Figure S9:**
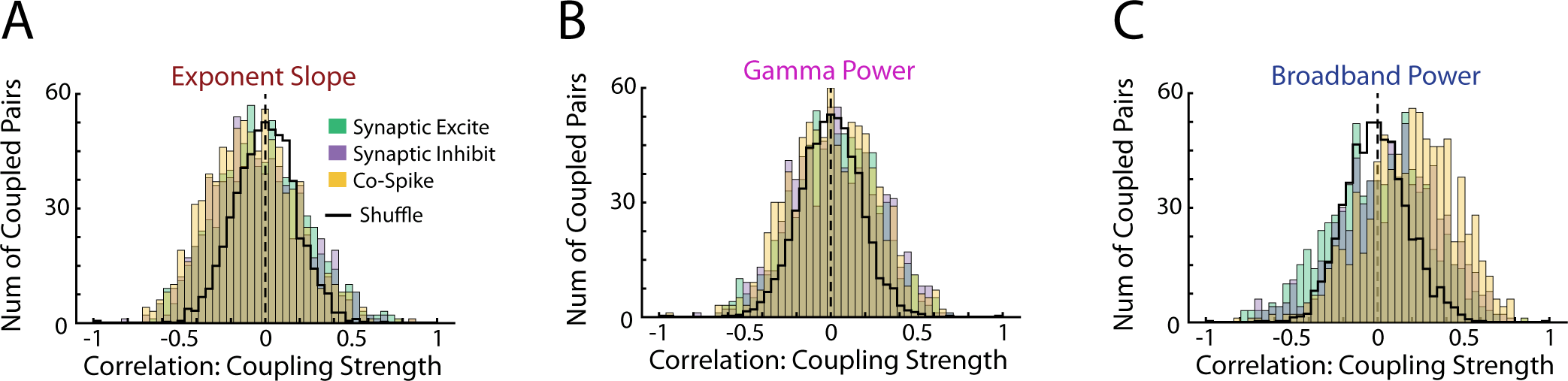
Relationship of co-spiking strength and LFP components. (A-C) Distribution of correlations between synaptic coupling or co-spiking strength and the LFP components of interest. Data are presented as in Fig 4A-C. Synaptic data are replotted from Fig 4 with co-spiking data now plotted for comparison purposes.

**Figure S10:**
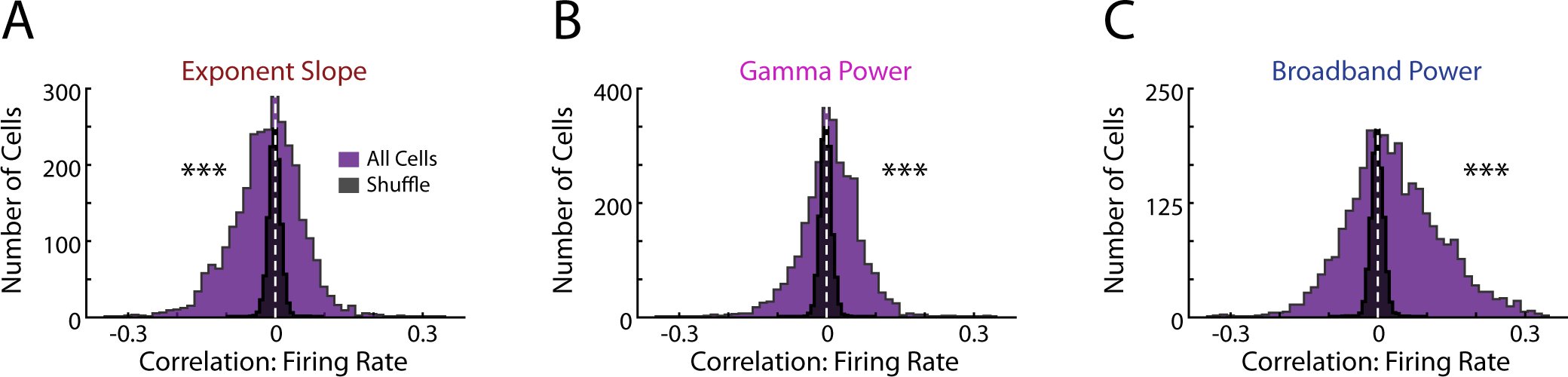
Correlation between mPFC firing rates and LFP components. (A-C) Distributions of correlations between the firing rates of all mPFC cells and the LFP components of interest. Data are presented as in Fig 4G-I but are no longer segregated according to principal cells and interneuron. Mann-Whitney U test vs shuffle; Exp. Slope: z=-9.46, p<0.001, d’=0.32; Gamma: z=11.29, p<0.001, d’=0.22; Broadband: z=17.01, p<0.001, d’=0.53. ANOVA across metrics: F(2)=349.40, p<0.001; post-hoc HSD Broadband vs Exp. Slop: p<0.001; Broadband vs Gamma: p<0.001.

**Figure S11:**
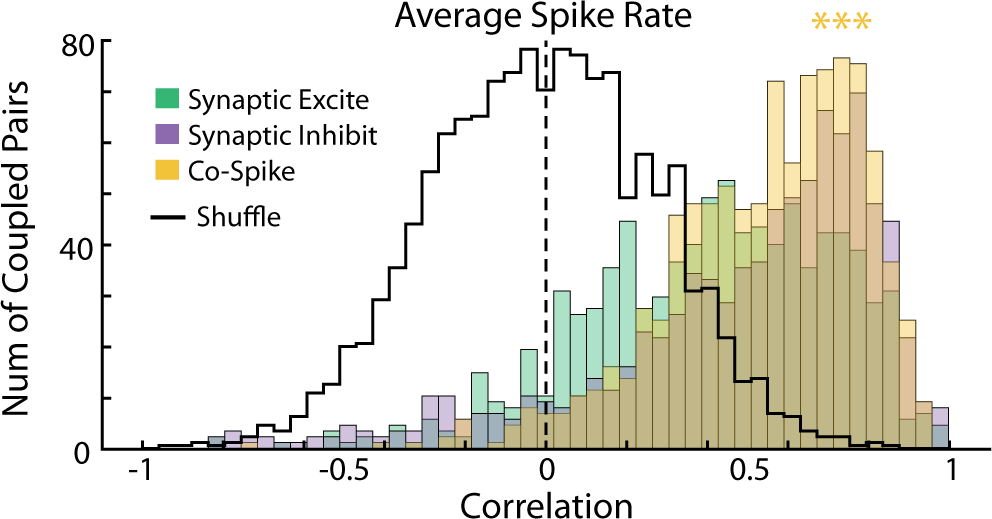
Relationship between firing rate and co-spiking strength. Distribution of correlations between the average firing rate of coupled mPFC cell pairs and the strength of the coupling relationship. Data are presented as in Fig 4J with synaptic data replotted and co-spiking data now included for comparison. Mann-Whitney U test; Co-Spike vs shuffle: z=31.04, p<0.001, d’=2.13.

